# Diversity of coelomycetous fungi in human infections: a 10-year experience of two European reference centres

**DOI:** 10.1101/323618

**Authors:** Dea Garcia-Hermoso, Nicomedes Valenzuela-Lopez, Olga Rivero-Menendez, Ana Alastruey-Izquierdo, Josep Guarro, José F. Cano-Lira, Alberto M. Stchigel, the French Mycoses Study Group

**Author notes:** Corresponding author; Unitat de Micologia, Facultat de 19 Medicina i Ciències de la Salut, IISPV, Universitat Rovira i Virgili, 21 Sant Llorenç St., 20 43201, Reus, Spain. **D.G-H. and N.V-L contributed equally to this article.**.

## Abstract

The coelomycetous fungi are difficult to properly identify from their phenotypic characterization and their role as etiologic agents of human infections is not clear. We studied the species distribution of these fungi among clinical isolates that had been collected and stored over a ten-year period in two European reference laboratories (France and Spain). We identified phenotypically and molecularly 97 isolates by sequencing the D1-D2 fragment of the 28S nrRNA (LSU) gene. Species of the orders *Pleosporales* and *Glomerellales* were present in both collections, and *Botryosphaeriales* and *Diaporthales* only in the French one. The most prevalent species were *Medicopsis romeroi*, *Neocucurbitaria keratinophila*, *Neocucurbitaria unguis-hominis* and *Paraconiothyrium cyclothyrioides*, which had been recovered primarily from superficial tissues. The *Didymellaceae* was the most common family represented, with 27 isolates distributed into five genera.

## INTRODUCTION

Human infections by coelomycetous fungi are rare and poorly characterized due to the difficulty in identifying these fungi using only phenotypic tools. The coelomycetous fungi are characterized by the production of conidia into fruiting bodies (= conidiomata), and were originally included in the orders *Sphaeropsidales* and *Melanconiales* of the class Coelomycetes, taxa which today lack scientific validity due to the demonstrated polyphyletic character of this sort of fungus (1–3). They cause superficial or subcutaneous infections, mostly following a traumatic inoculation of contaminated plant material or soil particles during agricultural work in tropical and subtropical areas (4–6). The most common coelomycetous fungi involved in these infections are the etiologic agents of black-grain eumycetoma, such as *Biatriospora mackinnonii; Falciformispora* spp., *Medicopsis romeroi*, and *Pseudochaetosphaeronema larense*. Other common coelomycetous fungi include *Lasiodiplodia theobromae* and *Neoscytalidium dimidiatum* (synanamorph of *Hendersonula toruloidea*) (7–11), which typically cause onychomycosis, subcutaneous phaeohyphomycosis (12–15), and eumycetoma (16). In addition, many species of *Phoma* and *Pyrenochaeta* have been reported as occasional agents of localized and systemic infections in humans (9, 17–20). The taxonomy of several coelomycetous genera mentioned before have been revised recently but they still constitute a group of highly polyphyletic taxa that are usually difficult to identify phenotypically (2, 21–24).

In a recent study conducted in the USA, Valenzuela-Lopez *et al.* (6) identified 230 fungal strains by sequencing the D1-D2 domains of the 28S rRNA gene (LSU), from which 152 (66.1%) strains belonged to the order *Pleosporales*, the rest being distributed in several orders of the phylum Ascomycota. Most of these strains were recovered from superficial tissue. *Neoscytalidium dimidiatum, Paraconiothyrium cyclothyrioides* and members of the family *Didymellaceae* were the most prevalent taxa. In addition, those authors demonstrated the usefulness of the LSU as a good molecular marker for a preliminary identification of coelomycetous fungi at genus level. In fact, such locus is easily amplified and many sequences are available in the GenBank database. However, the nucleotide sequences of more phylogenetically informative genes need analysing in order to identify the fungi at species level. Genes such as the RNA polymerase II subunit 2 (*rpb*2), translation elongation factor 1-alpha (*tef*1), beta-tubulin (*tub*2) and the ribosomal internal transcribed spacer region (ITS), combined in a multi-locus analysis, have all been recommended for this purpose (25)

Until now, the coelomycetous fungi involved in invasive fungal infections (IFIs) are poorly known in Europe, probably due to the infrequency of these fungi and the complexity of their identification in the absence of characteristic fruiting bodies when grown on culture media used in the clinical lab. In a recent French study, eighteen proven cases of cutaneous and subcutaneous primary infections by coelomycetous fungi were reported and analysed in patients from tropical and subtropical regions (26).

For a better knowledge of the diversity of coelomycetous fungi involved in human infections, we studied a large set of clinical isolates that had been identified in two mycology reference centres in France and Spain, and determined their *in vitro* antifungal susceptibility pattern.

## RESULTS

### Locations of infections

The majority of the isolates were recovered from superficial tissue, mainly skin (44%; 43/97), eyes (27%; 26/97), nails/hairs (18%; 17/97) and mouth/sinus (2%; 2/97). A few were recovered from deeper sites: bones (4%, 4/97), blood (2%, 2/97), cerebrospinal fluid (n=1), bone marrow (n=1) and lung (n=1) (Table 1 & 2).

**TABLE 1.**
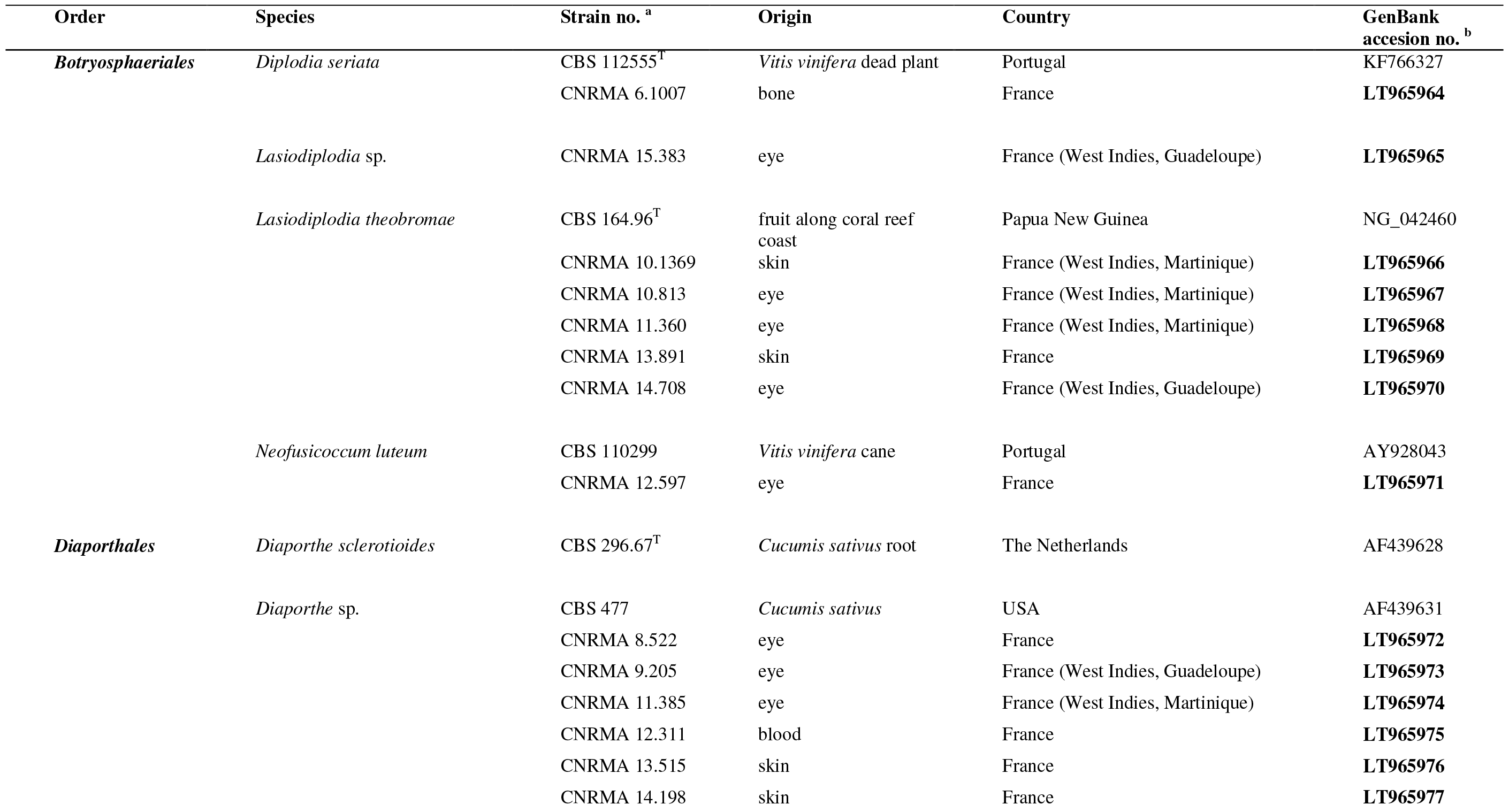
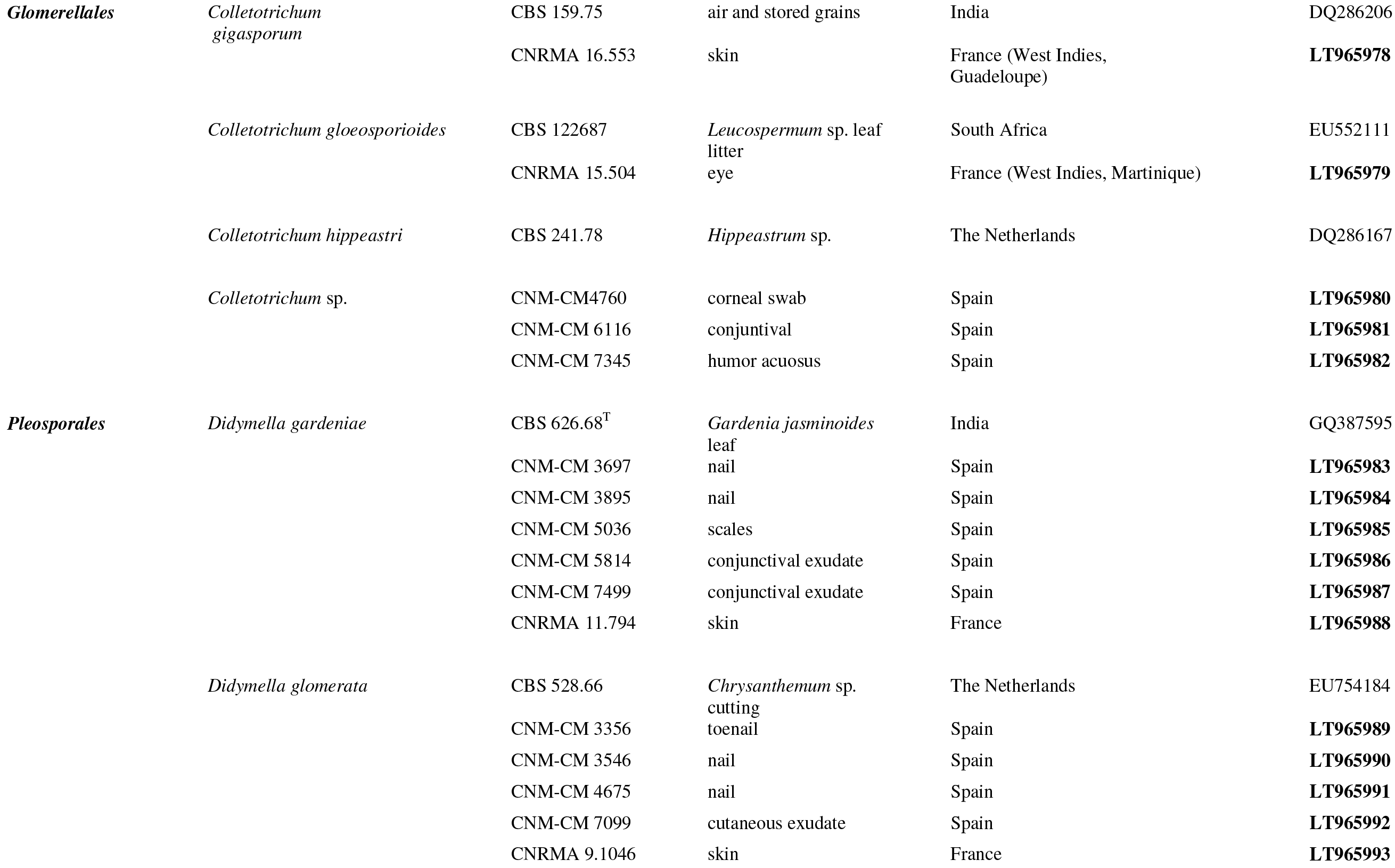
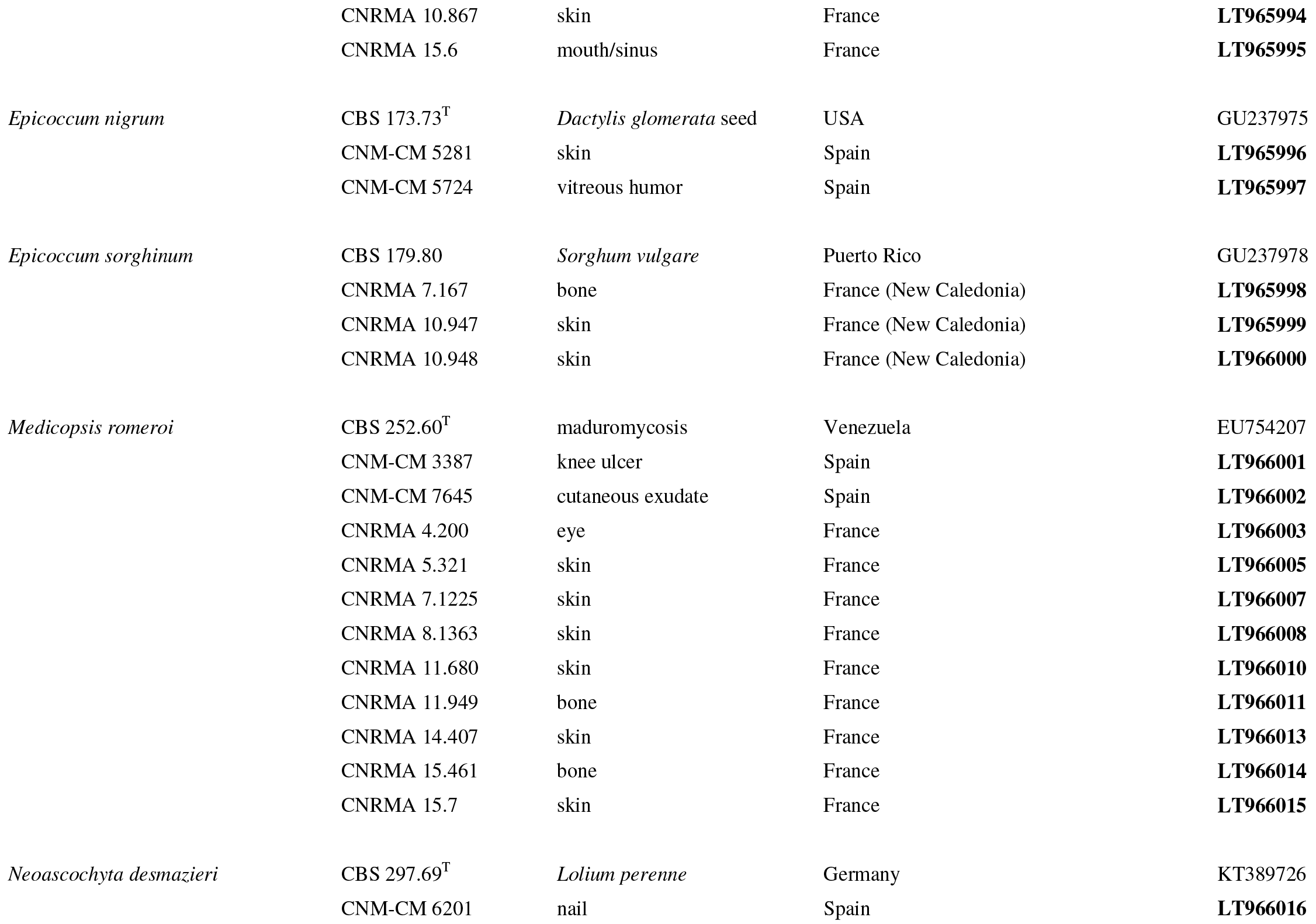
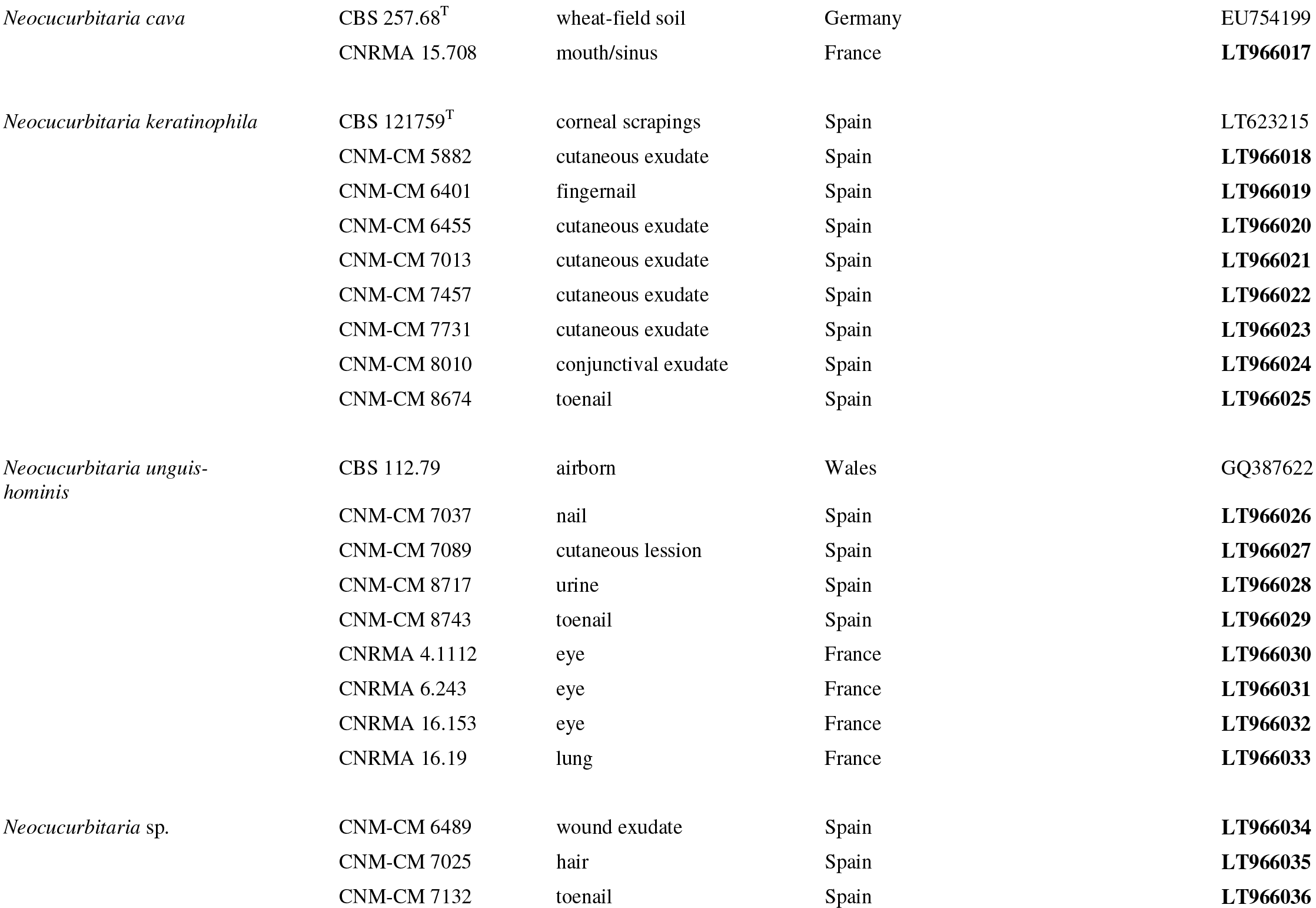
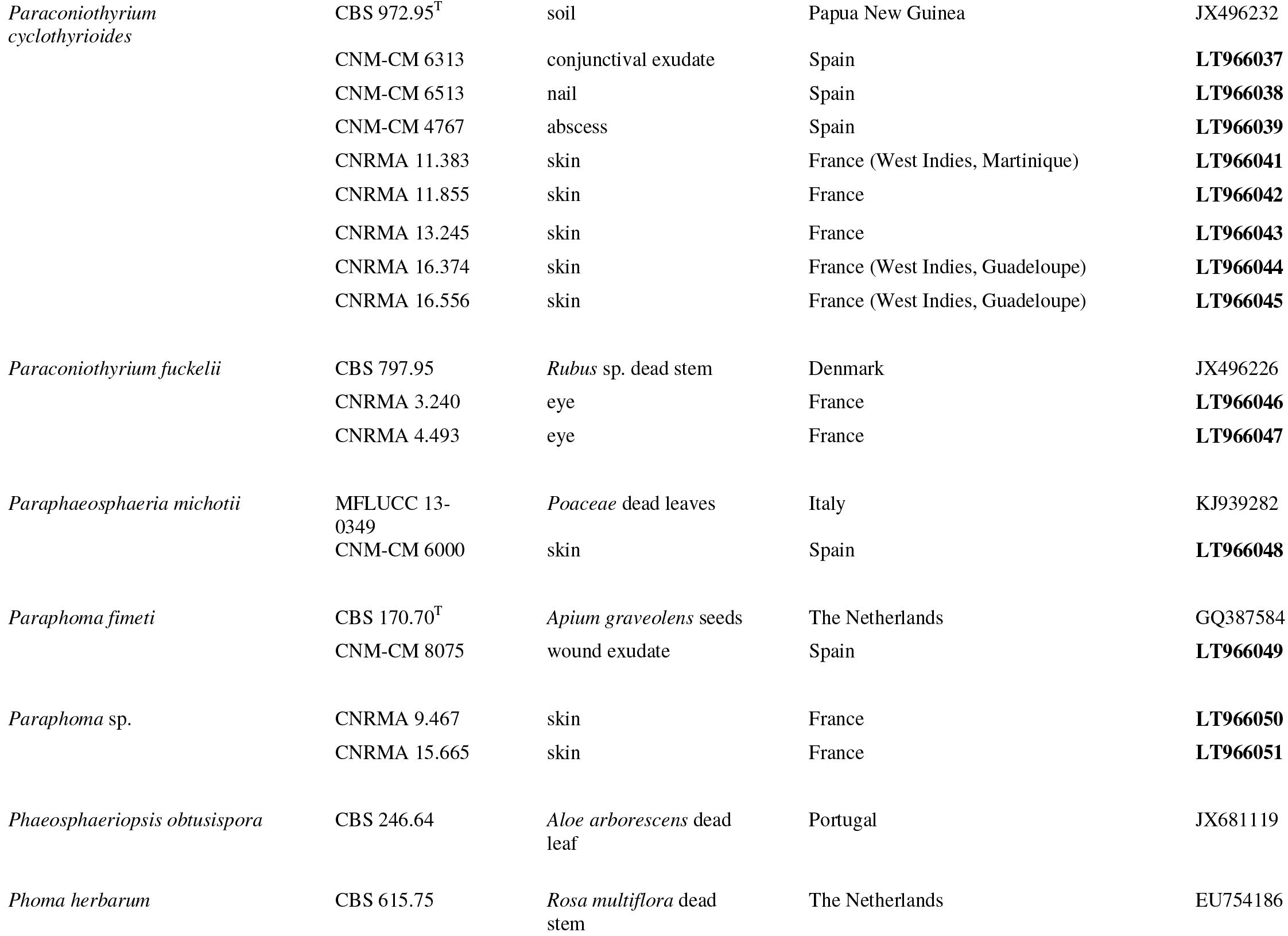
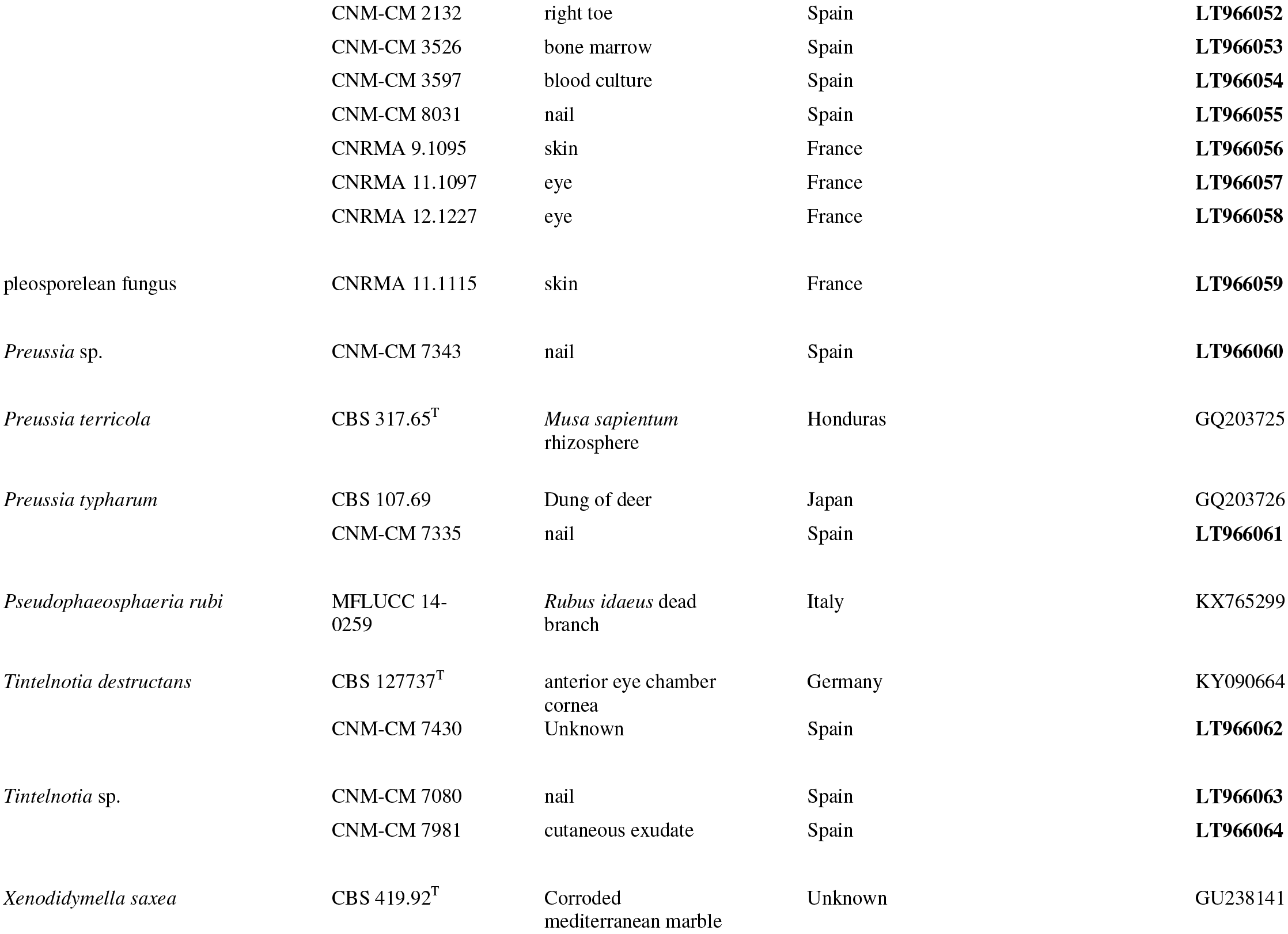
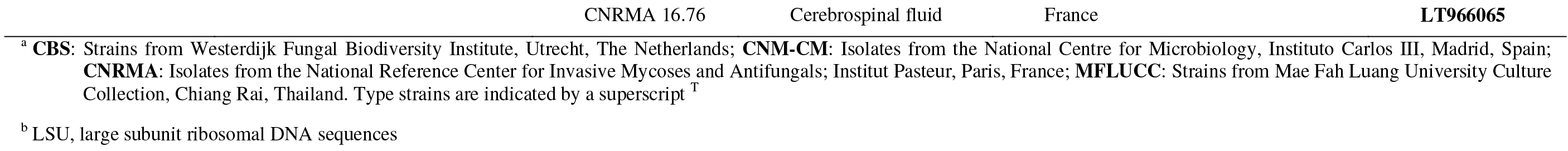
Taxonomical identification of the isolates studied, origin and GenBank accession numbers. New sequences generated are indicate in bold.

### Phylogenetic analyses

The maximum-likelihood (ML) phylogenetic analysis of the LSU sequences (approximately 584 pb) demonstrated that the 97 isolates were distributed into four orders, but scattered into fourteen clades (Fig. 1). Most of the isolates (81%; 78/97) belonged to the order *Pleosporales*, which were distributed into nine clades corresponding to 23 species of twelve genera, followed by those of the *Botryosphaeriales* (8%; 8/97), the *Diaporthales* (6%; 6/97) and the *Glomerellales* (5%; 5/97).

**FIG 1.**
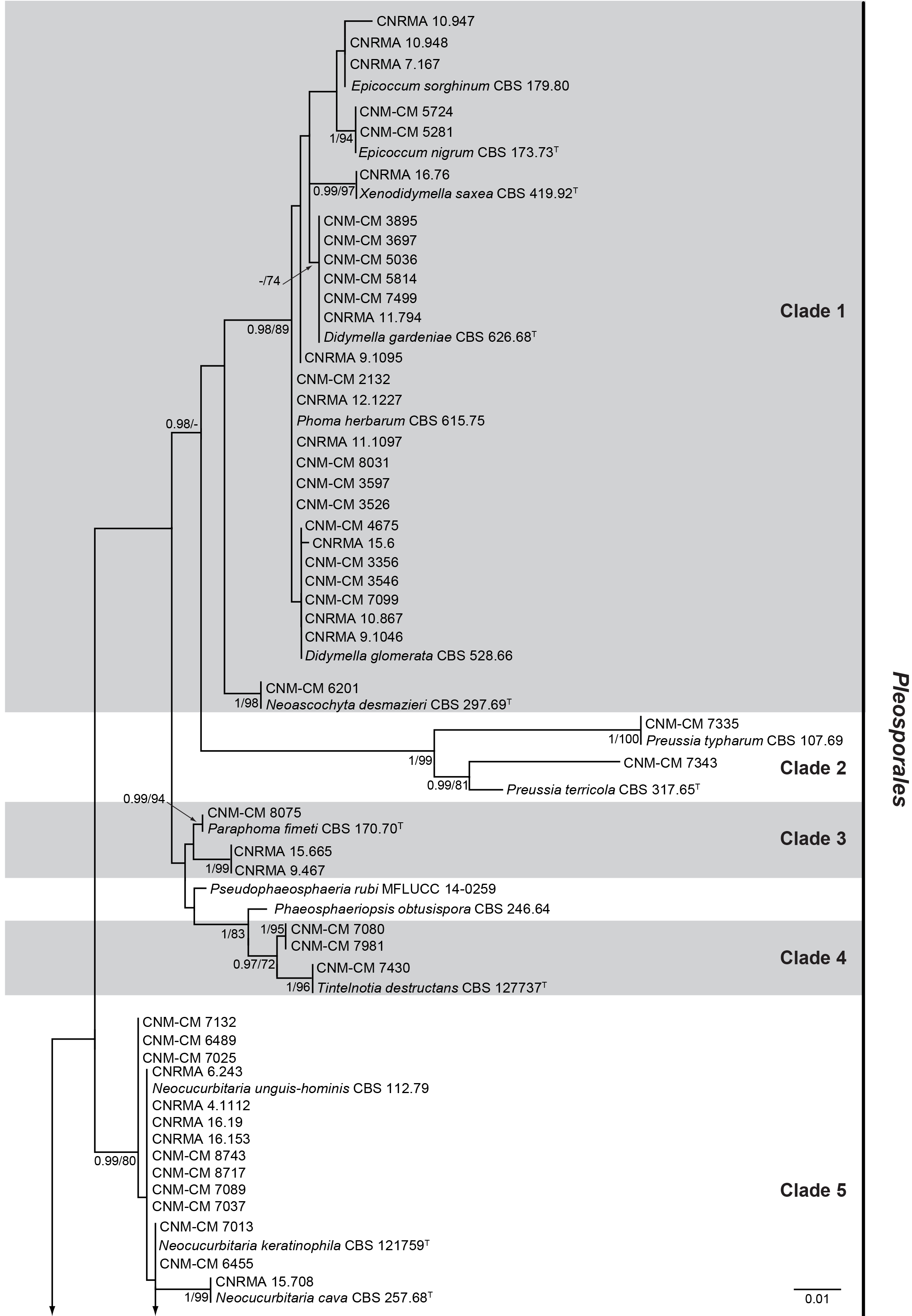
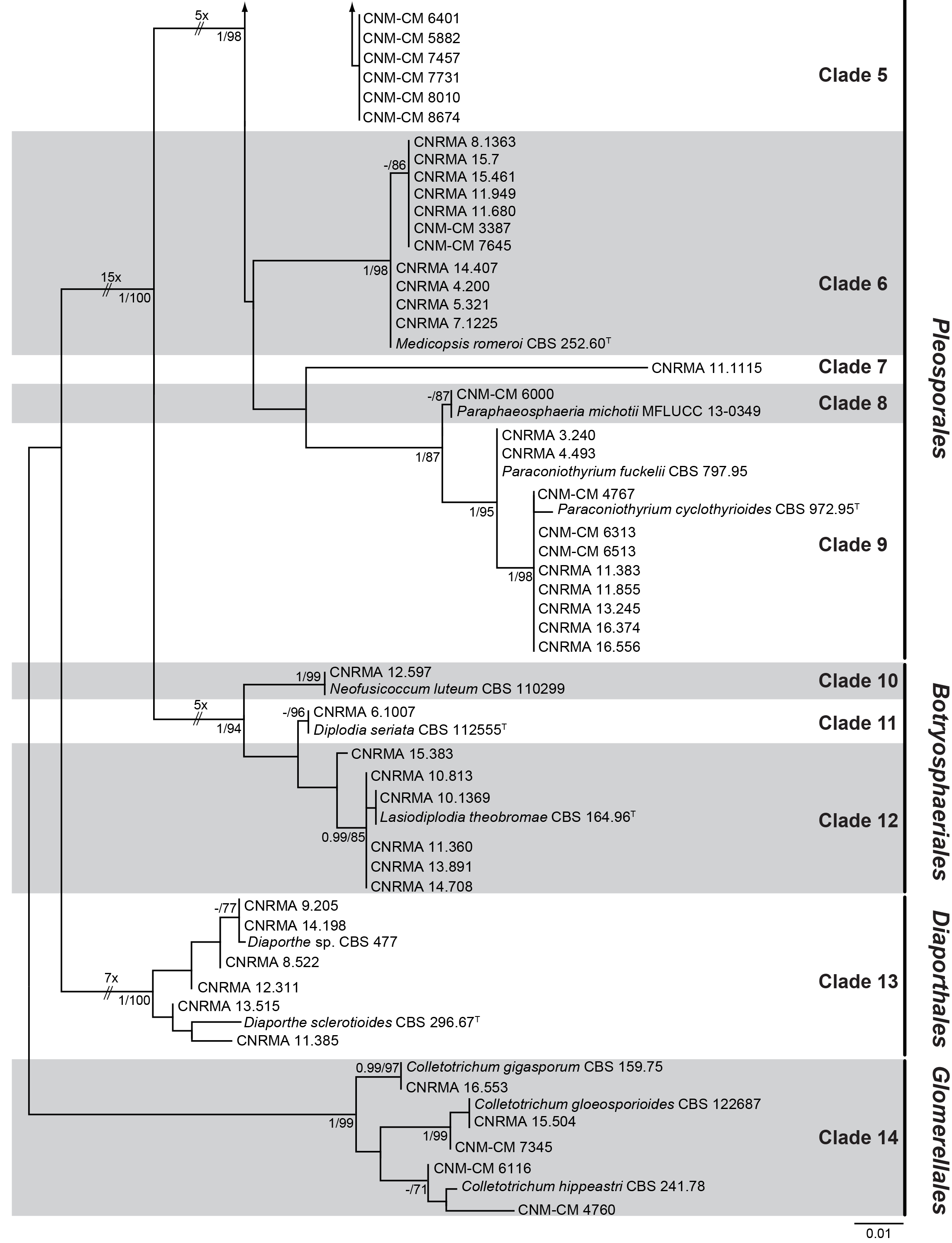
Maximum likelihood tree obtained from the D1-D2 of LSU (584 bp) sequences of the 125 strains, where 28 belong to type or reference strains. The branch lengths are proportional to phylogenetic distance. Bayesian posterior probability scores ≥ 0.95 and Bootstrap support values ≥ 70% are indicated on the nodes. Some branches were shortened to fit them to the page, these are indicated by two diagonal lines with the number of times a branch was shortened. The species of the genus *Colletotrichum* were used to root the tree. Superscript ^T^ indicated the type strains.

The most common species identified was *Medicopsis romeroi* (11%; 11/97), followed by *Paraconiothyrium cyclothyrioides*, *Neocucurbitaria keratinophila* and *N. unguis-hominis* (8% each; 8/97). These species were mostly isolated from cutaneous lesions (Table 2).

**TABLE 2.** Localization of infections due to coelomycetous fungi isolates

| **Orders** | no. of isolates obtained from: |  |  |
| --- | --- | --- | --- |
|  | **Superficial infection** | **Deep infection** | **Total no. of isolates** |
| *Botryosphaeriales* | 7 | 1 | 8 |
| *Diaporthales* | 5 | 1 | 6 |
| *Glomerellales* | 5 |  | 5 |
| *Pleosporales* | 71 | 7 | 78 |
| **Total no. of isolates (%)** | **88 (91)** | **9 (9)** | **97 (100)** |

Clade 1 of the *Pleosporales* corresponded to the family *Didymellaceae*, which included 27 isolates distributed into five genera, morphologically characterized by their production of pycnidial conidiomata and hyaline, aseptate conidia. The five genera were *Didymella*, *Epicoccum*, *Neoascochyta*, *Phoma* and *Xenodidymella*. *Didymella* was represented by 13 isolates, six of them clustering with the type strain of *D. gardeniae* (CBS 626.68), and the other seven clustered with a reference strain of *D*. *glomerata* (CBS 528.66). The genus *Epicoccum* grouped five of the isolates, three of them clustering with a reference strain of *E. sorghinum* (CBS 179.80) and the other two with the type strain of the type species of the genus, *E*. *nigrum* (CBS 173.73). The genus *Phoma* was represented by seven clinical isolates and a reference strain of *Phoma herbarum* (CBS 615.75). Two additional isolates included in this clade (CNRMA 16.76 and CNM-CM 6201) grouped with the type strains of *Xenodidymella saxea* (CBS 419.92) and *Neoascochyta desmazieri* (CBS 297.69), respectively.

Clade 2 had two species of *Preussia*: CNM-CM 7335 grouped with a reference strain of *P. typharum* (CBS 107.69), while CNM-CM 7343 represented an unknown species forming a sister clade with the type strain of *P. terricola* (CBS 317.65).

Clade 3 grouped three isolates of *Paraphoma*, one of them (CNM-CM 8075) clustered with the type strain of *P. fimeti* (CBS 170.70), and the remaining two (CNRMA 15.665 and CNRMA 9.467) representing unidentified phoma-like species.

Clade 4 had two sister clades of the genus *Tintelnotia*, which produced pycnidia and hyaline, aseptate conidia. The isolate CNM-CM 7430 was identified as *T. destructans*. However, the other two isolates (CNM-CM 7080 and CNM-CM 7981) did not cluster with any known species of the genus and might represent new species.

Clade 5 had 20 isolates of *Neocucurbitaria. Neocucurbitaria keratinophila* and *N. unguis-hominis* were the most common species, both with eight isolates each. *Neocucurbitaria cava*, with a single isolate (CNRMA 15.708), was also included in this clade. Three Spanish isolates, CNM-CM 6489, CNM-CM 7025 and CNM-CM 7132 were identified as *Neocucurbitaria* sp. due to being phylogenetically different from the other isolates and, again, might be a new species of the genus. *Neocucurbitaria* spp. produces pycnidia, ornamented or not, with bristle-like setose structures, and hyaline, aseptate conidia.

Clade 6 had eleven isolates of *Medicopsis romeroi* (syn. *Pyrenochaeta romeroi)*, which produces pycnidia and hyaline, aseptate conidia.

Clade 7 is represented by a single isolate (CNRMA 11.1115), phylogenetically distinct from the known pleosporalean fungi, possibly representing a novel taxon.

Clades 8 and 9 belonged to the family *Didymosphaeriaceae*. Clade 8 included a single isolate (CNM-CM 6000) phylogenetically related to a reference strain of *Paraphaeosphaeria michotii* (MFLUCC 13-0349). Clade 9 grouped ten isolates, two related to a reference strain of *Paraconiothyrium fuckelii* (CBS 797.95) and eight with the type strain of *Paraconiothyrium cyclothyrioides* (CBS 972.95). Members of the *Didymosphaeriaceae* form pycnidia and pale brown, 0-1 septate conidia.

The order *Botryosphaeriales* are present in Clades 10 to 12. Clade 10 had only one isolate (CNRMA 12.597) which clustered with a reference strain of *Neofusicoccum luteum* (CBS 110299); Clade 11 also had a single isolate (CNRMA 6.1007) that clustered with the type strain of *Diplodia seriata* (CBS 112555), and Clade 12 grouped six isolates, five of them clustering with the type strain of *Lasiodiplodia theobromae*, and CNRMA 15.383 identified as *Lasiodiplodia* sp. These fungi produce stromatic conidiomata and aseptate, hyaline to brown, thick-walled conidia.

Clade 13 included the type strain of *Diaporthe sclerotioides* (CBS 296.67) and six isolates corresponding to unidentified species of the genus *Diaporthe* (*Diaporthales*), none of them able to be morphologically distinguished since they produce pycnidia and small hyaline conidia.

Clade 14, corresponding to the *Glomerellales*, was used as outgroup. Five isolates nested in the *Colletotrichum* clade, two clustering with reference strains of *C. gigasporum* (CBS 159.75) and *C. gloeosporioides* (CBS 122687), respectively; and the other three, could not be identified. All the isolates showed the typical morphology of *Colletotrichum*, i.e., acervuli, conidia variable in shape, flattened with thickened tip branches (appressoria).

### Antifungal susceptibility testing

The minimum inhibitory concentration (MIC) was determined for 46 of the isolates included here (16 from Spain and 30 from France) (Table 3, Table S1). Globally, the geometric mean (GM) and MIC_50_ values of itraconazole and caspofungin were the highest (Table 3). The MIC of amphotericin B (0.06-1 mg/L) was generally low among the *Pleosporales* with the exception of one isolate of *M. romeroi* and one of *D. gardeniae*, with MICs of 8 and 32 mg/L, respectively. The azole MIC ranged between 0.03 and 1 mg/L for isolates belonging to the genera *Paraconiothyrium*, *Paraphoma*, *Tintelnotia* and *Neocucurbitaria*, with the exception of two isolates of *N. unguis-hominis*, which showed higher values (16 mg/L). The terbinafine MIC was low except for *Diaporthe* spp. and a few isolates of *Colletotrichum* spp. and *M. romeroi*.

**TABLE 3.** Overall in vitro antifungal activity against the 46 coelomycetous isolates as determined by EUCAST^a^ methodology

| Antifungal agent | MIC/MEC values (mg/L)<sup>b</sup> |  |  |  |  |
| --- | --- | --- | --- | --- | --- |
|  | range | median | GM | MIC<sub>50</sub> | MIC<sub>90</sub> |
| Amphotericin B | 0.03 -16 | 0.5 | 0.41 | 0.25 | 1 |
| Itraconazole | 0.014 -16 | 2 | 1.72 | 0.5 | 16 |
| Voriconazole | 0.03 -16 | 0.5 | 0.70 | 0.6 | 4 |
| Posaconazole | 0.014 -16 | 0.5 | 0.58 | 0.25 | 8 |
| Caspofungin | 0.125- 16 | 2 | 2.17 | 1 | 8 |
| Micafungin | 0.015- 16 | 0.5 | 0.53 | 0.125 | 8 |
| Terbinafin | 0.014- 16 | 0.25 | 0.39 | 0.25 | 2 |

<sup>a</sup>EUCAST, European Committee on Antimicrobial Susceptibility Testing procedure (71);
<sup>b</sup>MIC, minimum inhibitory concentration; MEC, minimal effective concentration; MIC<sub>50</sub> and MIC<sub>90</sub>, MIC encompassing 50 and 90% of isolates tested, respectively.

## DISCUSSION

The present study is the largest on this taxonomically complex group of fungi from clinical origin, with almost a hundred isolates morphologically and molecularly characterized from two southern European countries (France and Spain). Most of these coelomycetous fungi belonged to the order *Pleosporales* and were most commonly recovered from superficial infections. Similar results were observed in a previous work that focused on coelomycetous fungi collected at a North American reference centre (6). However, the diversity of the fungi identified in that study was higher, i.e. eleven orders were represented against four here.

In the present study, *Medicopsis romeroi* was the most frequently isolated species whereas the most common taxon in the American study was *Neoscytalidium dimidiatum*. Interestingly, while *M. romeroi* is usually reported as an etiologic agent of black grain eumycetoma (4, 11, 26–29), our isolates were mainly recovered from eye and non-mycetoma subcutaneous infections.

The second most frequently isolated species were *Paraconiothyrium cyclothyrioides*, *Neocucurbitaria unguis-hominis* and *N. keratinophila. Paraconiothyrium cyclothyrioides* is an emerging pathogen (6, 26, 30, 31) and was represented by eight isolates recovered from skin or superficial locations and mainly from tropical regions. *Neocucurbitaria unguis-hominis*, initially described as an agent of human onychomycosis (17), was equally distributed across both centres (n=8 isolates). Regarding *N. keratinophyla*, this species was reported for the first time from a corneal infection in Spain (18, 19). Interestingly, as well as being the first case reported for this species, all the isolates of *N. keratinophyla* were recovered in Spain from superficial tissue.

Other coelomycetous fungi we identified in the present work were *Didymella glomerata* and *Phoma herbarum*. Although *Phoma* spp. are commonly reported as a coelomycete involved in human infections (9, 20, 32–39), recent extensive changes in taxonomy and nomenclature have spread all but one of the species into different genera of the *Didymellaceae*, *Phoma herbarum* remaining as the unique species of the genus (22–24). Interestingly, *Didymella gardeniae* was commonly found in our study (five isolates from Spain and one from France).

Recently, Ahmed *et al.* (40) proposed *Tintelnotia destructans*, a new phoma-like fungus belonging to the *Phaeosphaeriaceae* able to cause eye and nail infections. They reported the successful use of terbinafine against a case of keratitis by this species. Two of the Spanish isolates recovered from superficial specimens (one cutaneous exudate and one nail sample) were molecularly related to the above-mentioned species but phylogenetically different and might represent a new taxon.

*Lasiodiplodia theobromae* (order *Botryosphaeriales*) is the only species of this genus involved in human opportunistic infections (41–46). Valenzuela-Lopez *et al.* (6) found a higher species diversity in the North American study than we report here, since five of the French isolates were identified as *L. theobromae*. The other three isolates of the *Botryosphaeriales* we found were related, one to a different species of *Lasiodiplodia* and the other two to other genera, specifically *Neofusicoccum* and *Diplodia*.

Four species of the genus *Diaporthe* (formerly *Phomopsis*; order *Diaporthales*), i.e. *D. bougainvilleicola*, *D. longicolla*, *D. phaseolorum* and *D. phoenicicola*, are considered opportunistic pathogens that cause mycoses that range from superficial to deep infections (47–51). Six isolates from France were phylogenetically placed into the latter genus. However, our results are only preliminary since only one phylogenetic marker was analysed. Similar was observed in several polyphyletic genera of the coelomycetes (52, 53).

We also report the finding of five clinical isolates of *Colletotrichum*. Two of the isolates corresponded to *C. gigasporum* (formerly *C. crassipes*) and *C. gloeosporioides*, taxa that have previously been reported as agents of keratitis, endophthalmitis and phaeohyphomicotic cyst; the other three isolates could not be identified at species level. This genus encompasses numerous plant pathogens that are found worldwide, although mainly in tropical and subtropical regions (54). The taxonomy of *Colletotrichum* is complicated and the genus is organized in species-complexes (55–59). Species such as *C. coccodes*, *C. crassipes*, *C. dematium*, *C. gloeosporioides*, *C. graminicola* and *C. truncatum* cause superficial and deep infections (endophthalmitis, keratitis, subcutaneous cyst or more rarely arthritis) (60–65). Further studies, including different phylogenetic markers, are needed to delimit the different species and clarify their pathogenic role.

The antifungal susceptibility of coelomycetous fungi involved in human infections is poorly known, mainly because they do not easily sporulate. In spite of the limited number of isolates tested here, amphotericin B seemed the most active drug *in vitro* together with terbinafine, in agreement with Valenzuela-Lopez *et al.* (6). Until more *in vitro* data is available, the antifungal treatment of the infection by this sort of fungus remains purely empirical. In a recent study, Guégan *et al.* (26) recommended extensive surgical resection of affected tissues as a first-line treatment for solitary subcutaneous lesions by coelomycetous fungi, followed by an antifungal therapy (posaconazole or voriconazole) in the case of relapse or amphotericin B in refractory cases.

Since our study is based on isolates from the two reference centres, we cannot comment on the incidence of infections due to coelomycetes nor compare their epidemiology between France and Spain. However, we still provide a good picture of the great diversity of coelomycetous fungi in the clinical context, and the basis for future studies on this interesting but neglected group of fungi.

## MATERIAL AND METHODS

### Fungal isolates

We studied 97 isolates of coelomycetous fungi recovered from clinical specimens, 51 of which were provided by the French National Reference Centre for Invasive Mycoses and Antifungals (NRCMA) at the *Institut Pasteur*, Paris (CNRMA isolates, n=51). The NRCMA offers expertise on difficult-to-identify fungi and the epidemiological surveillance of all cases of IFIs, which are notified on a voluntary basis either through active or passive surveillance programmes. The Spanish National Centre of Microbiology at the *Instituto de Salud Carlos III*, Madrid provided 46 isolates (CNM-CM isolates, n=46). This mycology reference laboratory receives isolates from the National Health System on a voluntary basis, the main aim of which is to support it by identifying and profiling the antifungal susceptibility of fungal isolates. The isolates were collected between 2005 and 2015. Table 1 gives information about the country of isolation and the location of the infection in the body.

### Morphological and physiological characterization

For morphology studies, the isolates were cultured on oatmeal agar (OA; 30 g of filtered oat flakes, 15 g of agar-agar, 1 L tap water) and malt extract agar (MEA; 40 g of malt extract, 15 g of agar-agar, 1 L distilled water) at 20 ± 1°C for 14 days in darkness. The morphological features of the vegetative and reproductive structures were studied using an Olympus CH2 bright-field microscope (Olympus Corporation, Tokyo, Japan) in wet mounts (on water and lactic acid) and slide cultures (by growing the isolates on OA and MEA) of the fungal isolates, following Valenzuela-Lopez *et al.* (6). Colour standards by Kornerup & Wanscher (66) were used in colony description. Photomicrographs were taken with an Axio-Imager M1 microscope (Zeiss, Oberkochen, Germany).

### DNA extraction, amplification and sequencing

Total genomic DNA was extracted from colonies grown on potato dextrose agar (PDA; 4 g of potato infusion, 20 g dextrose, 15 g of agar-agar, 1 L tap water) after seven days of incubation at 20 ± 1°C, using the FastDNA kit protocol (Bio101, Vista, CA), with a FastPrep FP120 instrument (Thermo Savant, Holbrook, NY) following the manufacturer’s protocol. DNA was quantified using the Nanodrop 2000 (Thermo Scientific, Madrid, Spain). LSU was amplified with the primer pair LR0R and LR5 (67). The amplicons were sequenced in both directions with the same primer pair used for amplification at Macrogen Europe (Macrogen Inc., Amsterdam, The Netherlands). The consensus sequences were obtained using the SeqMan software version 7.0.0 (DNAStar Lasergene, Madison, WI, USA).

### Molecular identification and phylogenetic analysis

Preliminary molecular identification of the isolates was made using LSU nucleotide sequences in BLAST_N_ searches. Twenty-eight LSU sequences of type or reference strains deposited in the GenBank database by the Westerdijk Fungal Biodiversity Institute (CBS) and the Mae Fah Luang University (MFLUCC) culture collections were used for identification and phylogenetic purposes. DNA sequences generated in this study were deposited in GenBank (accession numbers are given in Table 1).

For the phylogenetic study, sequences were aligned using the ClustalW application (68) of the MEGA 6.06 (69) computer program, and manually adjusted using the same software platform. Phylogenetic reconstructions were made by maximum-likelihood (ML) and Bayesian inference (BI) with MEGA 6.06 and MrBayes 3.2.4 (70), respectively. The best substitution model for the gene matrix (TN93+G) was estimated using MEGA 6.06. For ML analyses, nearest-neighbour interchange was used as the heuristic method for tree inference. Support for internal branches was assessed by 1,000 ML bootstrapped pseudoreplicates. Bootstrap support (BS) of ≥70 was considered significant. For BI analyses, Markov chain Monte Carlo (MCMC) sampling was carried out with four million generations, with samples taken every 1,000 generations. The 50% majority rule consensus trees and posterior probability values (PP) were calculated after removing the first 25% of the resulting trees for burn-in. A PP value of ≥0.95 was considered significant. Reference strains of *Colletotrichum gigasporum* (CBS 159.75), *C. gloeosporioides* (CBS 122687) and *C. hippeastri* (CBS 241.78) were used as outgroup.

### Antifungal susceptibility testing

The *in vitro* susceptibility testing in both reference centres (n= 46 isolates) followed the European Committee on Antimicrobial Susceptibility Testing (EUCAST) procedure (71, 72). The antifungals used were amphotericin B (Sigma-Aldrich Química, Madrid, Spain), itraconazole (Sigma-Aldrich Química, Madrid, Spain), posaconazole (Schering-Plough Research Institute, Kenilworth, N.J.), voriconazole (Pfizer S.A., Madrid, Spain), caspofungin (Merck & Co., Inc., Rahway, N.J.), micafungin (Astellas Pharma Inc, Tokyo, Japan) and terbinafine (Novartis, Basel, Switzerland). For the NCRMA, all antifungal drugs were obtained from ALSACHIM, Strasbourg, France.

The isolates were cultured on potato carrot agar (PCA; 20 g each of filtered potatoes and carrots, 20 g of agar, 1 L of distilled water) or OA for seven to 30 days at 25°C and 30°C to obtain sporulation. Conidia were then collected in sterile water containing 0.01% (v/v) Tween 80 (Sigma-Aldrich, St. Louis, MO, USA), and the suspension was adjusted to 2–5 × 10^5^ conidia/mL. The minimal effective concentration (MEC) was determined for each echinocandin and the minimal inhibitory concentration (MIC) for the other drugs (90% inhibition for amphotericin B and 80% for the azoles) after 24 h and 48 h of incubation at 35°C. *Aspergillus flavus* ATCC 204304 and *Aspergillus fumigatus* ATCC 204305 were used as quality control strains in all tests carried out. Susceptibility profiles were determined for 46 isolates since non-sporulating isolates were excluded at the NRCMA.

## ACKNOWLEDGMENTS

We thank Cécile Gautier (National Reference Center for invasive Mycoses and Antifungals [NRCMA]) for technical assistance. This work was supported by the Spanish *Ministerio de Economía y Competitividad*, grant CGL2017-88094-P.

Olga Rivero-Menendez holds a fellowship from the *Fondo de Investigaciones Sanitarias* (grant FI14CIII/00025). Ana Alastruey-Izquierdo is supported by a research project from the *Fondo de Investigación Sanitaria* (FIS) (project PI16CIII/00035).

**Members of the French Mycoses Study Group** who contributed their clinical isolates to this study are as follows: Annecy (S. Bland), Bichat (C. Chochillon), Boulogne-Billancourt (N.Ait-Ammar), Caen (C. Duhamel), Clermont Ferrand (P.Poirier), Cochin (A. Paugam), Corbeil-Essones (D. Kubab), Guadeloupe (M. Nicolas), La Réunion (S. Picot), Lyon (A.L. Bienvenu), Martinique (N. Desbois), Nantes (F. Morio), Necker (M.E. Bougnoux), Nouvelle Calédonie ( R. Goursaud), Pitié (A. Fekkar), Poitiers (C. Kauffmann), Quinze-Vingts (L. Merabet), Rouen (L. Favennec), Saint Etienne (H. Raberin), Saint-Louis (A. Alanio), Toulouse (S. Cassaing).

